# Design to Data for mutants of β-glucosidase B from *Paenibacillus polymyxa*: I45K, A357S, I20A, I20V, and I20E

**DOI:** 10.1101/2020.10.07.330233

**Authors:** Jennie A. Luong, Ashley Vater, Justin B. Siegel

## Abstract

The relatively small size and scope of most current datasets of biophysical mutation effects in enzymes limit the ability to develop data-driven algorithms enabling accurate generative modeling tools for designing novel enzyme function. Here, the Michaelis-Menten constants (*k*_cat_, K_M_, and *k*_cat_/K_M_) and thermal stability (T_M_) of five new mutations of β-glucosidase B from *Paenibacillus polymyxa* (BglB) are characterized. Foldit software was used to create molecular models of the mutants, for which synthetic genes were constructed and the corresponding proteins produced and purified from *E. coli*. It was found that mutations that disrupted pre-existing hydrogen bonds near the active site had reduced expression in contrast to mutations at the same site that did not affect native hydrogen bonding. This is consistent with previous results showing the relationship between hydrogen bonding and enzyme functionality. These mutants contribute to a growing data set of >100 mutants that have been characterized for expression, kinetic, and thermal properties

## INTRODUCTION

Enzymes are the primary means of catalyzing chemical reactions in biological systems and are becoming more important in the development of therapeutics.^1,2^ Protein engineering can be used to create stable, efficient enzymes for reactions that are not catalyzed by natural biocatalysts.^1^ Two factors that are often key considerations when creating biocatalyst are (1) the denaturation temperature and (2) the efficiency of reaction catalyzation.^3^ While large standardized datasets for protein structures exist,^4^ the field lacks large, quantitative standardized datasets of enzyme functional properties. While powerful and accurate tools for modeling protein structure are rapidly evolving, the lack of data has limited the ability to develop robust predictive algorithms for a proteins functional properties.^5,6^ While structure is critical for function large and consistently generated data sets are needed to better capture the subtleties from which a proteins structure imparts its function.^7^

To address this gap there are growing efforts to create large data sets of large data sets directly connecting protein sequence-structure-function relationships. Of particular relevance to this study are efforts to quantitatively characterize a growing library of mutants for the *Paenibacillus polymyxa beta-glyhcosidase B* (BglB), for which >100 mutants have now been produced and systematically characterized for expression, kinetic, and thermal properties.^8,9^ In this study, a total of five single-point BglB mutants were modeled, synthetic genes constructed, and corresponding proteins biophysically characterized. Mutants were designed using Foldit^10^. The thermal stability of the mutant was predicted by the total system energy score (TSE)^6^ and catalytic efficiency was predicted by the ligand energy. Within this set, two of the mutant’s models showed a change in intermolecular forces and three preserved those of the native structure. The mutants A357S and I20E were predicted to have a decreased overall catalytic efficiency (*k*_cat_/K_M_) and mutants I20A, I20V, and I45K were predicted to have no change in overall catalytic efficiency. Mutant I45K was hypothesized to have an increase in overall thermal stability and all other mutants were predicted to have a decreased thermal stability. Based on the TSE and ligand energy alone, only three of the five mutants’ functional performance aligned with what was initially hypothesized. In this small sample set, there appears to be a trend between the variants’ expression, thermal stability, and catalytic efficiency in relationship to changes in the modeled intermolecular interactions—where small, non-interruptive changes had little effect on functionalities and disruptive changes considerably diminished functionality.

## METHODS

### Mutation Design

Foldit was used with a previously described method^8^ to model the five-point mutations. The mutations were scored by the Rosetta energy function and both a TSE and ligand energy were given.^6^ Mutants were chosen, in part, to have similar TSE values as the wild type, within 5 points of the wild type energy score.

### Generating Synthetic Genes of Mutants

Plasmids containing a synthetic gene encoding the mutant proteins were generated with standard Kunkel Mutagenesis procedure.^12^ Plasmids were electroporated into Dh5alpha competent *E*.*coli* cells and plated onto Luria-Bertani agar plates with 50 μg/mL kanamycin. Individual colonies were sequenced via sanger sequencing after overnight growth to identify mutants.

### Protein Purification and Production

The sequence verified BglB mutants were grown, and expressed using previously described methods.^9^ After inducing protein expression with IPTG, the cells were lysed and immobilized metal ion affinity chromatography was used to purify proteins. Protein yield was determined by A280 using a BioTek® Epoch spectrophotometer and protein purity was assessed by SDS-PAGE gel.

### Michaelis Menten Kinetics and Thermal Stability

Kinetic characterization was done using a previously described method.^8^ Activity of the enzyme was measured using the production rate of 4-nitrophenol from p-nitrophenyl-beta-D-glucoside. The measurements were determined and recorded through A420 spectrophotometry assay for 1 hour. *k*_cat_ and K_M_ were determined by fitting the data to the Michaelis-Menten kinetics model.^13^ The thermal stability (T_M_) for the variants were determined using Protein Thermal Shift™ kit made by Applied BioSystem ® from ThermoFisher. Following the standard protocol by the manufacturer, purified proteins were diluted from 0.1 to 0.5 mg/mL and fluorescence reading was monitored using QuantaStudio™ 3 System from 20 °C to 90 °C. The T_M_ values were then determined using the two-state Boltzmann model from the Protein Thermal Shift™ Software by Applied BioSystem ® from Thermo Fisher.

## RESULTS

### Interpreting Foldit Models

The mutants A357S and I20E were predicted to have a decreased *k*_cat_/K_M_ due to the positive change in ligand energy (Table 1), as well as observable changes in the hydrogen bonding adjacent to the active site, destabilizing the area (Figure 2). Mutants I20A, I20V, and I45K were predicted to have no change in *k*_cat_/K_M_ despite having a positive change in ligand energy (Table 1) because of the lack of observable changes to the intermolecular interactions between the substrate and the ligand (Figure 2). Mutant I45K was hypothesized to have an increase in overall thermal stability due to the negative change in the TSE value (Table 1). All other mutants were predicted to have a decreased thermal stability. From the data, we can see that mutants I20E, I45K, and A357S were predicted accurately for *k*_cat_/K_M_, while mutants I20A, I20V, and A357S were predicted accurately for thermal stability.

**Table 1.** Δ Total System Energy Score, Expression, *k*_cat_, K_M_, *k*_cat_/K_M_, T_M_ of five mutations and two wildtypes. Summary of the data collected for all mutants. The two wild type data sets have been combined as the average between the two. Technical triplicate data was collected for each mutant and all parameters, the standard deviation for the best fit data is provided.

| Mutants | $\Delta TSE$ | $\Delta LigandE$ | Yield (mg/mL) | $k_{cat}$ ($\text{min}^{-1}$) | $K_M$ (mM) | $k_{cat}/K_M$ ($\text{min}^{-1} \text{mM}^{-1}$) | $T_M$ ($^{\circ}\text{C}$) |
| --- | --- | --- | --- | --- | --- | --- | --- |
| WT (Average) | 0.000 | 0.000 | 0.523 | $622.0 \pm 45.8$ | $9.8 \pm 2.2$ | $62.9 \pm 11.5$ | $45.1 \pm 0.1$ |
| A357S | 0.117 | 0.237 | 0.148 | $92.8 \pm 3.2$ | $12.7 \pm 1.4$ | $7.3 \pm 0.8$ | $41.4 \pm 0.4$ |
| I45K | -3.148 | 0.009 | 1.136 | $299.9 \pm 2.4$ | $4.2 \pm 0.1$ | $72.0 \pm 2.4$ | $43.6 \pm 0.3$ |
| I20A | 1.693 | 0.742 | 0.168 | $92.7 \pm 2.1$ | $17.1 \pm 1.1$ | $5.4 \pm 0.4$ | $42.2 \pm 0.8$ |
| I20E | 1.133 | 0.300 | 0.145 | N/A | N/A | N/A | N/A |
| I20V | 0.381 | 0.090 | 1.225 | $564.0 \pm 7.5$ | $12.7 \pm 0.53$ | $44.5 \pm 1.9$ | $44.8 \pm 0.2$ |

**Figure 1.**
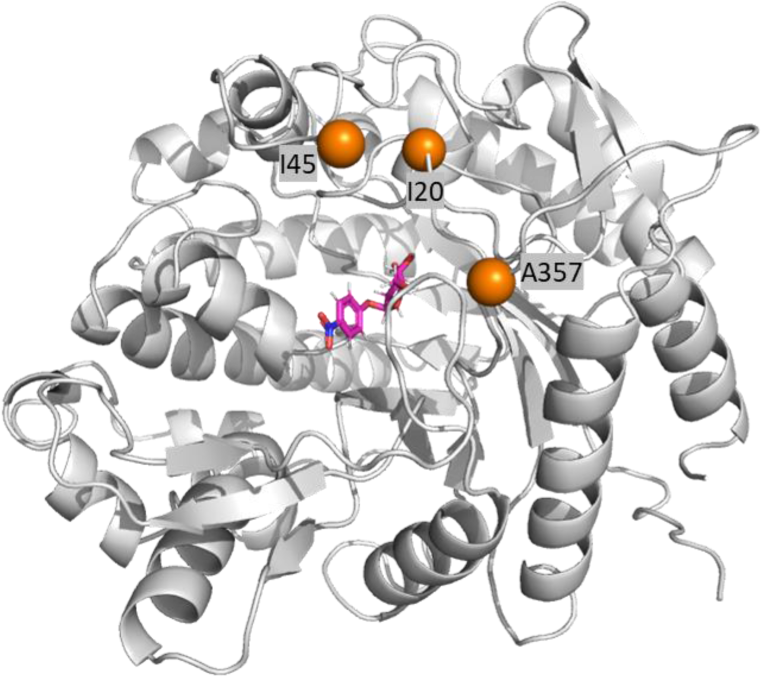
Location of mutant sites on BglB from Paenibacillus polymyxa. Pymol^11^ rendering of BglB. The three mutational sites used in the study are indicated in orange spheres. The substrate is indicated in pink and the main enzyme is shown in grey.

**Figure 2.**
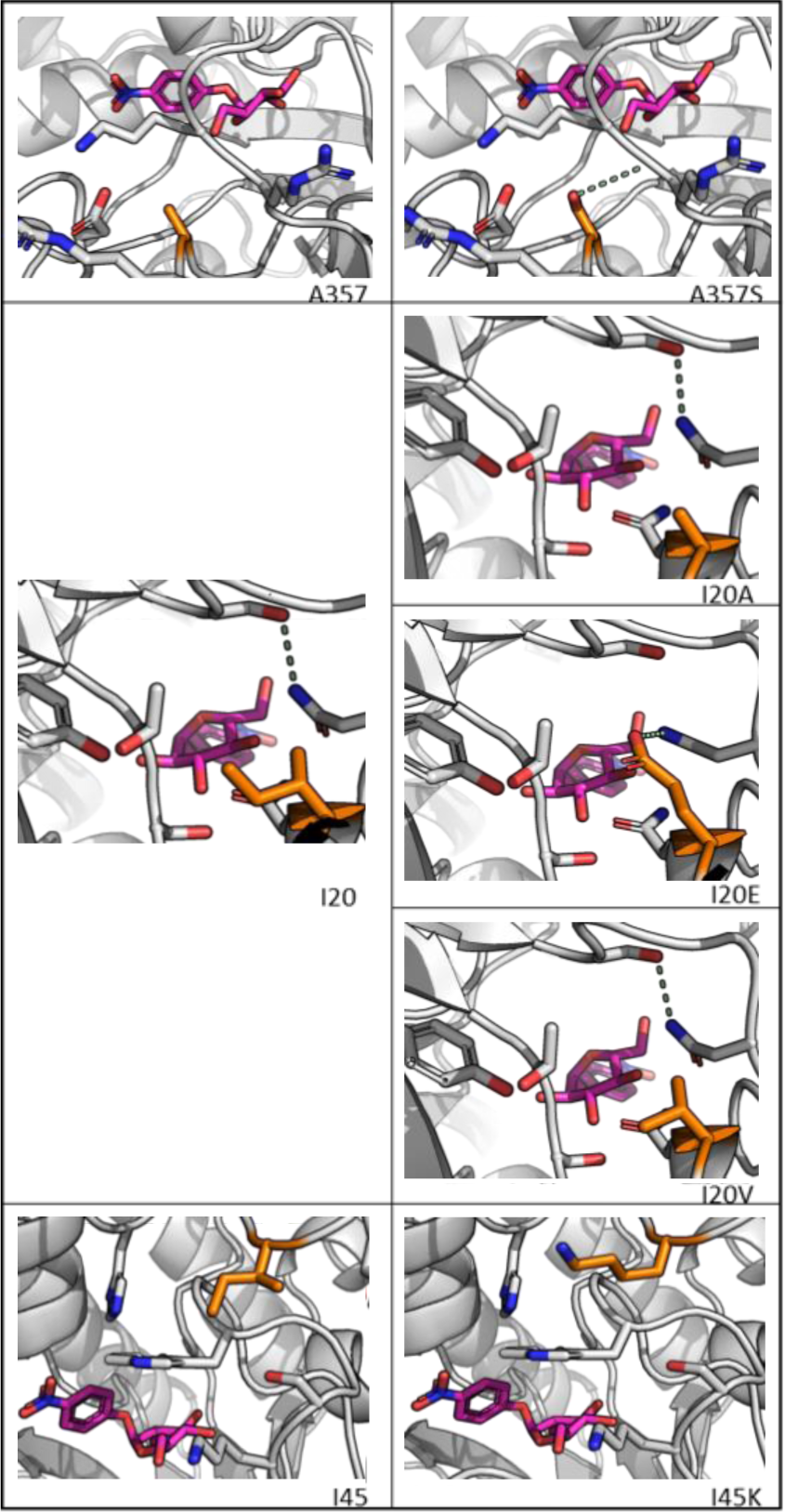
Structural Representation of BglB Mutants. Mutations are illustrated using PyMOL^6^ generated images showing the positions of the mutated amino acid (orange), the ligand (pink), and hydrogen bonds (green).

### Expression Results

All mutants expressed and purified as soluble protein above the detection limit (0.1 mg/mL) as previously defined in this system^6^ based on A280 and SDS-PAGE analysis. There were varying levels of expression with mutants A357S, I20E, and I20A, having the lowest yields, only slightly higher than the detection limit (Table 1) and the lightest bands (Figure 3). Mutants I45K and I20V expressed at a higher level, with yields >1.0 mg/mL concentration (Table 1) and the darkest bands (Figure 3).

**Figure 3.**
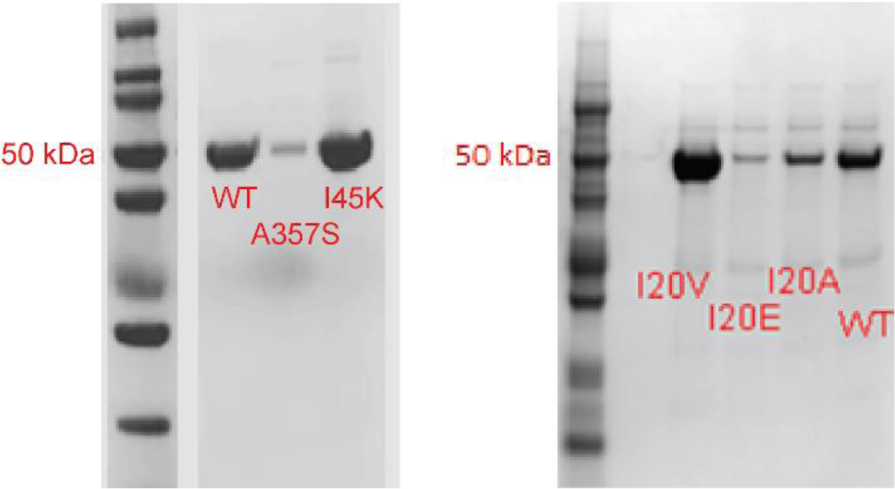
SDS-PAGE gel of two wild type and mutant enzymes. Sample bands formed at 50 kD, showing BglB purity. Mutants shown from left to right: I20V, 120E, 120A, A357S, 145K.

### Kinetic Results

The average wild type BglB *k*_cat_ / K_M_ value was 62.9 mM^-1^ min^-1^ (Table 1), which falls within the range of typical *k*_cat_/K_M_ values established by studies of this system ^8,9^ and across enzyme systems more broadly.^14^ While a protein band was present in the I20E lane on the SDS-PAGE analysis, the enzyme had no quantifiable activity. Using the previously established limit for enzymatic activity detection of 10 M^-1^min^-1 8^, only mutant (I20E) fell below this value. Mutant A357S had a K_M_ double that of the wildtype and a 77% drop in *k*_cat_ when compared to the wildtype, suggesting that interactions with the substrate were inhibited (Figure 4). For the mutants I20A and I20V, we saw decreases in overall catalytic efficiency (approximately 91% and 27%, respectively). Mutant I45K saw similar results to the wildtype in catalytic efficiency *k*_cat_/K_M_.

**Figure 4.**
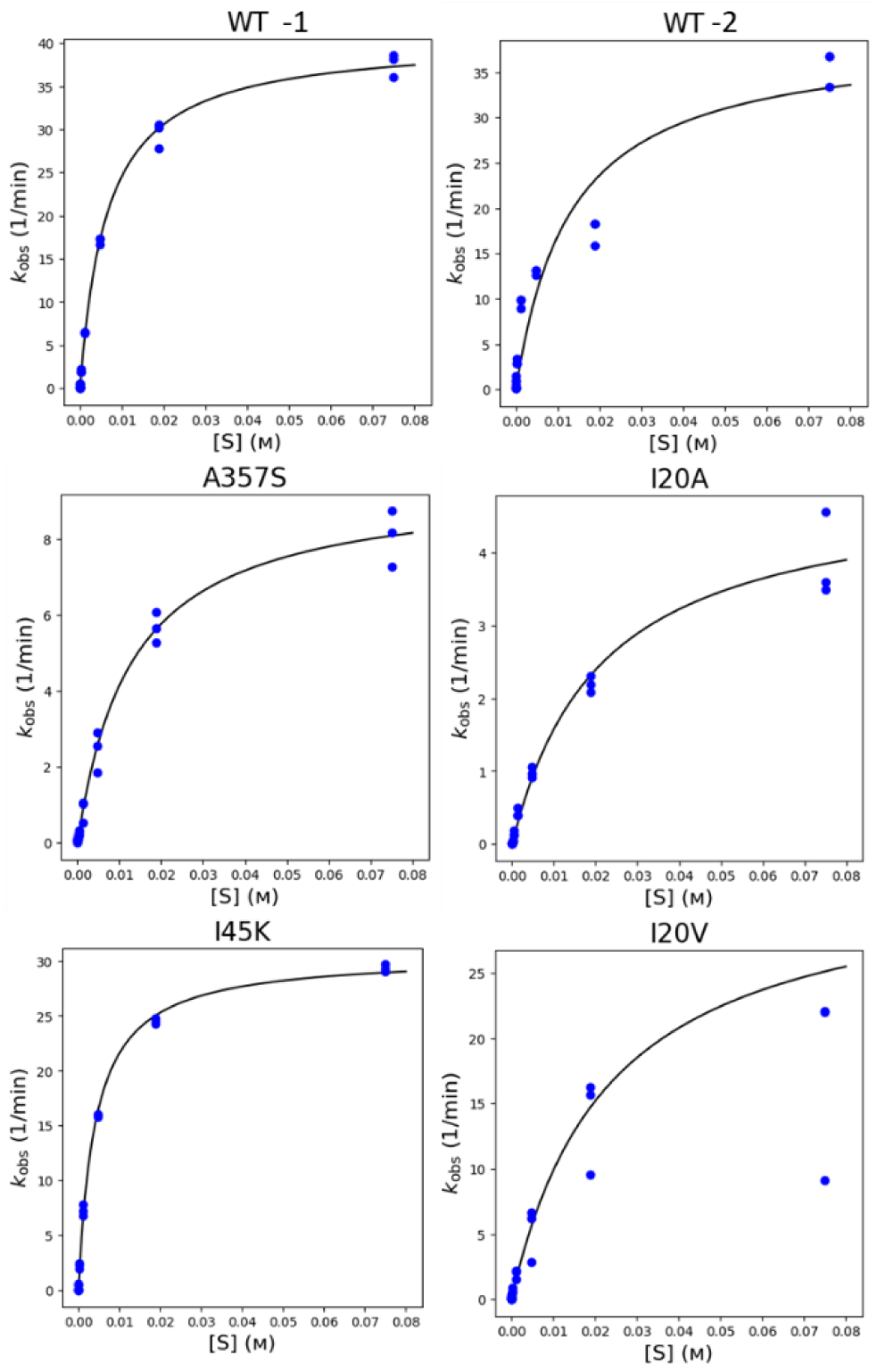
Observed turnover rate verses substrate concentration. Biological replicates of WT are presented as WT-1 and WT-2, all mutants and WT enzymes were assayed as technical triplicates with all data points shown in the plots. Concentrations of pNG are plotted on the horizontal axis with units of M and kobs on the vertical axis of min^-1^. The data above was fit through non-linear regression to the Michaelis-Menten equation to determine *k*_cat_ and K_M_, the best fit line for the triplicate data is illustrated in each panel.

### Thermal Stability Results

The mean T_M_ of wildtype BglB was determined to be 45.1 °C (Table 1). No thermal data was collected for mutant I20E due low protein yield making reading unreliable. All other mutants had a decrease in thermal stability. The T_M_ of the mutants A357S, I20A, I20V, and I45K were decrease by approximately 3 °C, 3 °C, 0.3°C, and 2 °C, respectively. All thermal data can be seen in Figure 5, as well as in Table 1.

**Figure 5.**
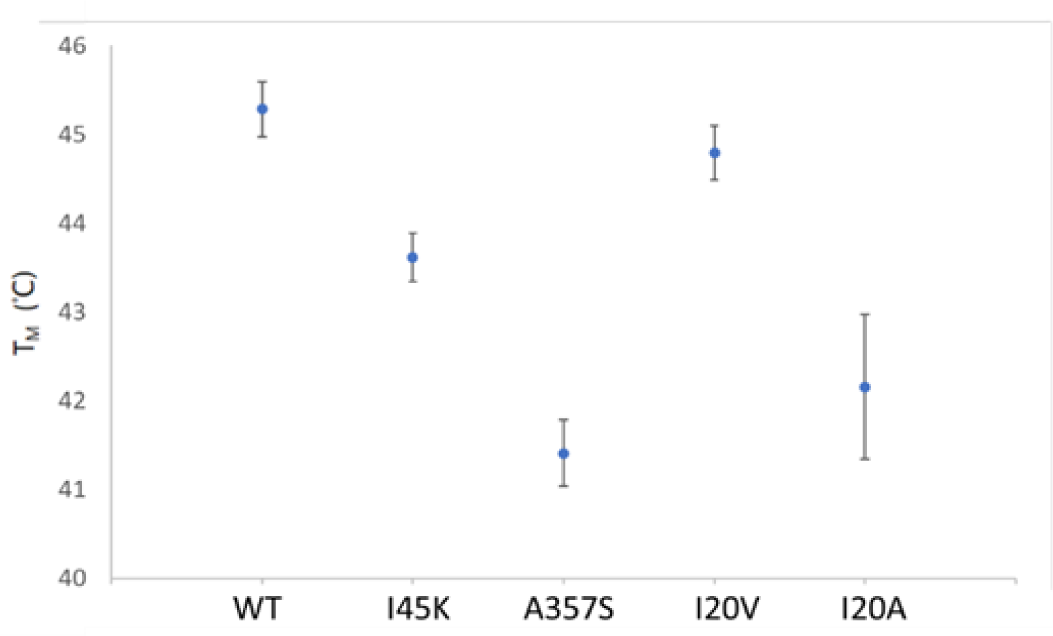
Thermal Stability of BglB Mutants. The T_M_ values are in units of degrees Celsius. Bars indicate standard error from triplicate samples. WT-1 and WT-2 replicates are combined and reported as one WT.

## DISCUSSION

In this study we found that three of the five predicted mutational effects for catalytic efficiency and three of the five predicted effects for thermal stability were supported by the data. Further, some patterns can be seen within the data that may be interesting to investigate.

Loss of catalytic efficiency was nearly ubiquitous across our small sample set; in reflecting back on the structural models, we think we may be seeing an effect from changes in hydrogen bonding near the active site and from changes in the size of hydrophobic interactions. For example, the complete loss of activity of I20E might be due to a loss of a hydrogen bond near the active site between S400 and N404 in exchange of a hydrogen bond between the mutant and N404. And the striking decrease in activity of A357S may have been due to the hydrogen bond that was created between the serine and the backbone of the protein structure, causing an unfavorable structural change for binding affinity. For both I20A and I20V, we hypothesized that the reduction we saw in catalytic efficiency might be due to the decrease in size of the hydrophobic group, resulting in a loss of hydrophobic interaction which decreased protein stability and substrate affinity. Similar results were previously observed with the mutant V401A in a study by Chengrui Hou et al 2019.^15^

To further support the idea that hydrogen bonding has a considerable effect, the mutant I45K did not seem to affect catalytic efficiency *k*_cat_/K_M_. This is may have been due to the lack of new hydrogen bonding that was formed, resulting in very similar interactions between the mutant and the wildtype.

We observed an overall trend between expression and activity and thermal stability. As expression (Yield, Table 1) increases, so does thermal stability and catalytic efficiency. Those that had the lowest levels of expression were the most disruptive mutations, causing changes to the hydrogen bonding that occurred *within* the protein structure (I20E and A357S) as opposed to changes occurring between the protein and the ligand. These are consistent with existing rules of protein folding.^16^

We can see this with more clarity when looking at the mutations at the I20 position. When comparing mutants I20A, I20V, and the wildtype, we think we see a relationship in that the size of a non-polar amino acid, defined by the number of carbon atoms in the amino acid, leading to a decrease in both thermal stability and catalytic efficiency. These mutants do not change the hydrogen bonding in the protein, in contrast to mutant I20E, where a hydrogen bond is lost in favor of another. This change in hydrogen bonding coincided with low expression levels and catalytic efficiency falling below the detectable level.

The effects that hydrogen bonding on enzyme functionality have been previously examined. It was observed that the hydrogen bond energy of protein side chains had the strongest correlation with the T_M_ of the enzyme when compared to other features.^9^ This was also observed in the Pearson correlations presented in Carlin et al. 2016, where there was a positive correlation between 1/K_M_ and the total number of hydrogen bonds in BglB.^8^ These findings align with the values for mutations at the I20 position in this study.

By focusing in on one site of the protein, we were able to observe how different interactions changed enzyme functionality. This study further stresses the importance of hydrogen bonding, not only between the substrate and the enzyme, but also within the structure of the enzyme. Finally, these findings show that focusing in on one position of a protein may help glean new information on enzyme functionality that may be useful in protein design. These mutants add to the Design to Data efforts to catalog the effects of a single point mutation on BglB.

## ACKNOWLEDGEMENTS

This work was supported by the University of California Davis, the National Institutes of Health (R01 GM 076324-11), the National Science Foundation (award nos. 1827246, 1805510, and 1627539), and the National Institute of Environmental Health Sciences of the National Institutes of Health (award no. P42ES004699). The content is solely the responsibility of the authors and does not necessarily represent the official views of the National Institutes of Health, National Institute of Environmental Health Sciences, National Science Foundation, or UC Davis.

